# Temperature adaptation of yeast phospholipid molecular species at the acyl chain positional level

**DOI:** 10.1101/2024.04.04.588069

**Authors:** Celine Kelso, Alan T. Maccarone, Anton I.P.M. de Kroon, Todd W. Mitchell, Mike F. Renne

## Abstract

The budding yeast *Saccharomyces cerevisiae* is a poikilothermic organism and adapts its lipid composition to the environmental temperature to maintain membrane physical properties. Studies addressing temperature-dependent adaptation of the lipidome in yeast have described changes in the phospholipid composition at the level of sum composition (*e.g.* PC 32:1) and molecular composition (e.g. PC 16:0_16:1). However, to date, there is no information at the level of positional isomers (*e.g.* PC 16:0/16:1 *versus* PC 16:1/16:0). In this study, combined Collision- and Ozone-Induced Dissociation (CID/OzID) mass spectrometry was deployed to investigate homeoviscous adaptation of PC, PE, and PS *sn*-molecular species composition. We determined the main species to be 16:1/16:1, 16:0/16:1, 16:1/18:1, 16:0/18:1, and 18:0/16:1. In general, at higher culture temperature, the *sn*-1 position is increased in saturated acyl chains, whereas the *sn*-2 position mainly is increased in acyl chain length. PC mainly increases in 16:0/16:1 and 16:0/18:1, at the expense of 16:1/16:1, whereas PS and PE increase in 16:1/18:1, at the expense of 16:1/16:1 and 16:0/16:1. Our data suggest distinct adaptation mechanisms of the *sn*-1 and *sn*-2 acyl chains, and different manners of *sn*-molecular species adaptation between PC and PE/PS.

## Introduction

The biophysical properties of biological membranes, such as the fluidity (viscosity), thickness, compressibility and permeability of the membranes, are highly dependent on the composition of membrane lipids (Harayama and Antonny 2023; Renne and Ernst 2023; Harayama and Riezman 2018; van Meer, Voelker, and Feigenson 2008). Glycerophospholipids (or phospholipids in short, PL) comprise the bulk of the membrane lipids, and therefore are critical for determining membrane properties.

The physico-chemical properties of membrane phospholipids depends on (*i*) the phospholipid class, which is defined by the nature of the lipid headgroup esterified to the phosphate group at the *sn*-3 position, and (*ii*) the PL molecular species, defined by the acyl chains linked to the *sn*-1 and *sn*-2 positions of the glycerol backbone (Renne and de Kroon 2018; van Meer, Voelker, and Feigenson 2008). To maintain membrane homeostasis, cells must monitor membrane physical properties and tightly regulate lipid metabolism (Los and Murata 2004; de Mendoza and Pilon 2019; Ernst, Ballweg, and Levental 2018; Ernst, Ejsing, and Antonny 2016). How cells perform this task at the molecular level, and how membrane lipid composition is altered to preserve membrane homeostasis, are fundamental questions in membrane biology.

The budding yeast *Saccharomyces cerevisiae* has been used extensively as a model system to study lipid metabolism (Santos and Riezman 2012; Singh 2016), and the metabolic pathways for PL biosynthesis have been well characterized (Figure 1A) (Henry, Kohlwein, and Carman 2012). Compared to higher eukaryotes, yeast has a simple acyl chain composition consisting of predominantly 16 or 18 carbon atoms, which are either mono-unsaturated or fully saturated (Martin *et al*, 2007; Wagner & Paltauf, 1994). The acyl chain composition is mainly determined by *de* n*ovo* fatty acid synthesis, followed by acyl-CoA desaturation and elongation (Tehlivets *et al*, 2007; De Kroon *et al*, 2013). Despite this simple acyl chain composition, the yeast lipidome comprises hundreds of structurally distinct lipids (Ejsing et al. 2009; Klose et al. 2012; Danne-Rasche, Rubenzucker, and Ahrends 2020; Reinhard et al. 2020). Phospholipids are the major component of the yeast lipidome, with phosphatidylcholine (PC), phosphatidylethanolamine (PE) and phosphatidylinositol (PI) being the most abundant classes (Ejsing et al. 2009; Klose et al. 2012).

**Figure 1.**
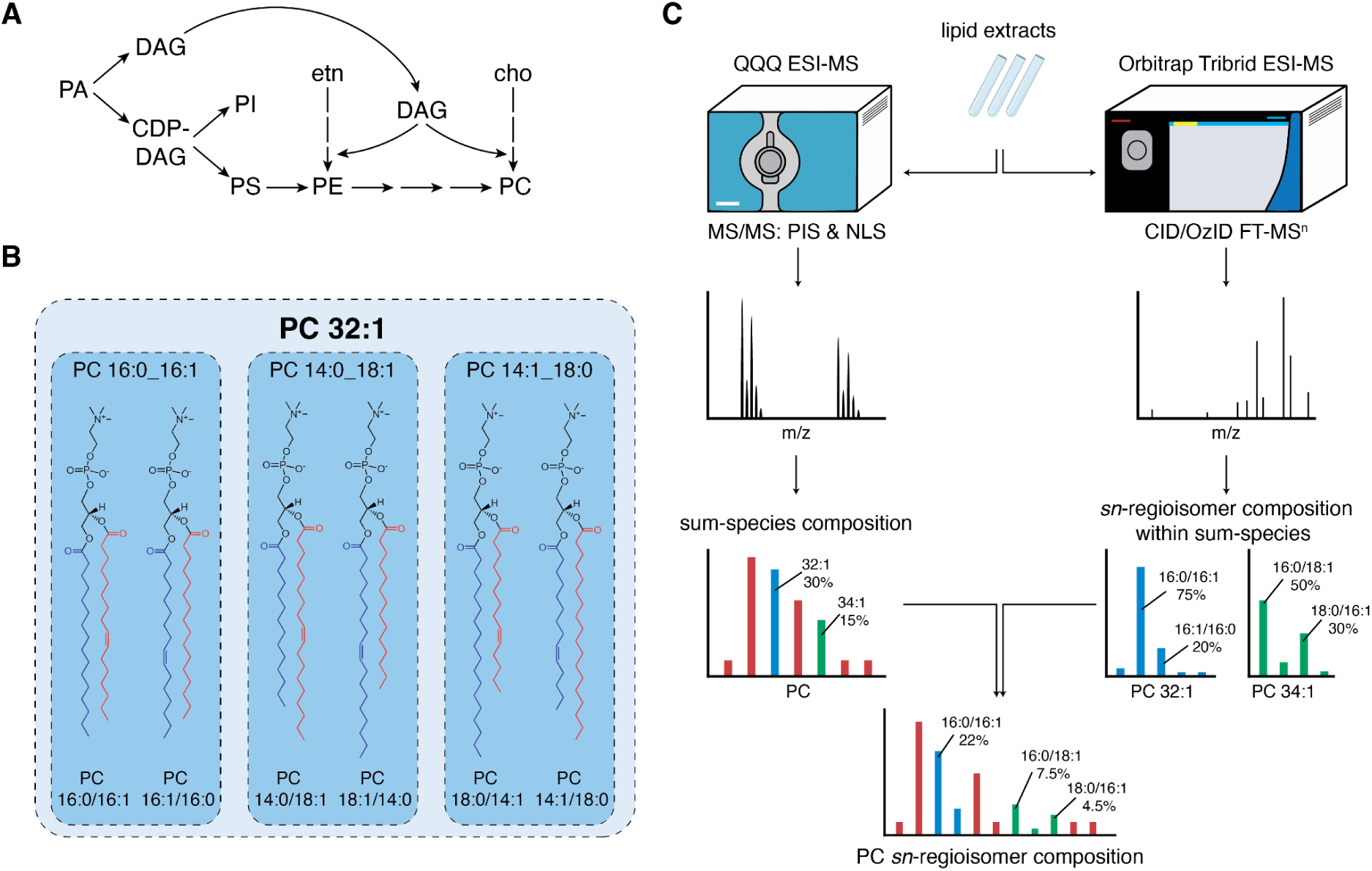
– Different levels of phospholipid identification. (A) Schematic overview of the biosynthetic pathways producing the major glycerophospholipids PC, PE, PS, and PI in yeast. (B) Phosphatidylcholine identified at the sum species level as PC 32:1, consists of acyl chains with a total of 32 carbon atoms, and one unsaturation. PC 32:1 can consist of multiple distinct species at the acyl chain level, identified by the two acyl chains esterified to the glycerol backbone, such as PC 16:0_16:1, PC 14:0_18:1 and PC 14:1_18:0 where the underscore indicates the absence of information on the sn-localization of the esterified acyl chains. Not taking into account the double bond location, each of these species has two sn-regioisomers, for example PC 16:0/16:1 (phosphatidylcholine with 16:0 at sn-1 and 16:1 at sn-2) versus PC 16:1/16:0. For further discussion on the lipid structural identification, the reader is referred to (Triebl et al. 2017; Hancock et al. 2017) (C) Schematic overview of 2 ESI-MS/MS-based workflows for lipid structural characterisation: traditional Neutral Loss & Precursor Ion scans on a triple quadrupole platform (left) and novel Ozone-Induced Dissociation-rooted methods coupled with accurate mass spectrometry on a modified Orbitrap (right).

In the last 3 decades, the yeast phospholipid composition has been investigated at various levels of lipid identification (Figure 1B). Tandem mass spectrometry approaches using precursor ion scanning and/or multiple reaction monitoring allowed the initial analysis of membrane phospholipid species at the sum-species level (e.g., PC 32:1) (Boumann et al. 2003; Guan and Wenk 2006; Schneiter et al. 1999). Shotgun lipidomics approaches combining high resolution mass spectrometry with MS^n^ analysis increased structural resolution and identified the lipid molecular composition at the level of fatty acyl chains esterified to the glycerol backbone (Ejsing et al. 2009; Reinhard et al. 2020). In line with the acyl chain composition of yeast, the main PC species were found to be 16:1_16:1, 16:0_16:1, 16:0_18:1, and 16:1_18:1 (Ejsing et al. 2009). However, this data does not provide the location of the esterified acyl chains, which defines the phospholipid positional isomers or *sn*-molecular species (*e.g.* distinguishing the *sn*-regioisomers PC 16:0/16:1 and PC 16:1/16:0). Selective hydrolysis of the *sn-1* or *sn-2* esterified acyl chains using specific phospholipases has been employed to examine the acyl chain composition at both positions. In a seminal work, Wagner & Paltauf showed that the phospholipid *sn*-2 position generally is enriched in unsaturated acyl chains, compared to the *sn*-1 position (Wagner and Paltauf 1994). Furthermore, the *sn-*1 position is enriched in C16 acyl chains whereas the *sn*-2 contains more C18 acyl chains (Wagner and Paltauf 1994). However, to date there seems to be little information on the yeast lipidome defined at the level of *sn*-molecular species.

Yeast is a poikilothermic organism and adapts its lipid composition to the environmental temperature to maintain membrane physical properties (Ernst, Ejsing, and Antonny 2016; Hazel 1997). This process was initially hypothesised to be driven by maintaining membrane viscosity, thus the process was named “homeoviscous adaptation” (Hazel 1997). Recent works indicate that membrane homeostasis goes beyond maintenance of membrane viscosity/fluidity (Renne and Ernst 2023; Harayama and Antonny 2023), yet it remains clear that cells must maintain membrane physical properties in response to changes in the environment. The first evidence for yeast lipidome adaptation in response to temperature established redistribution of the total acyl chain composition, increasing in acyl chain length and decreasing unsaturation in response to higher culture temperatures (Martin, Oh, and Jiang 2007; Suutari, Liukkonen, and Laakso 1990; Hunter and Rose 1972). Using shotgun lipidomics, Klose *et al*. showed that at elevated temperatures, phospholipids generally increase in monounsaturated species at the expense of diunsaturated species, and decrease in C32 species compensated by an increase in C34 species (Klose et al. 2012). Ejsing *et al*. used MS^2^ data to determine the phospholipid molecular composition, showing that with increasing temperatures, PC mainly increases in PC 16:0_16:1, PC 16:1_18:1, and PC 16:0_18:1 at the expense of PC 16:1_16:1 (Ejsing et al. 2009). Pursuant to the alterations in the PC molecular composition, PC increased in C16:0, C18:0 and C18:1 at the expense of C16:1. In addition to adaptation of the molecular species composition, the PL class composition also adapts to altered culture temperature, showing an increase in PC and PI at the expense of PE at higher temperatures (Ejsing et al. 2009; Klose et al. 2012).

Advances in analytical approaches have enabled the near complete structural characterization of lipids (Ekroos et al. 2003; Groessl, Graf, and Knochenmuss 2015; Pham et al. 2014; Maccarone et al. 2014; Triebl et al. 2017). Mass spectrometry-based lipid analysis using ozone-induced dissociation (OzID) allows for the identification of phospholipid species at the acyl chain position level, *i.e.* determining the acyl chains esterified to the *sn-*1 and *sn-*2 positions of the glycerol backbone (Pham et al. 2014; Poad et al. 2018; Hancock et al. 2017; Michael et al. 2024). Here, we employed collision-combined with ozone-induced dissociation mass spectrometry (CID/OzID) to determine the *sn*-molecular species composition of the aminoglycerophospholipids PC, PE, and phosphatidylserine (PS) of yeast cultured at different temperatures. Combining the CID/OzID data with the sum-molecular species composition obtained by electrospray ionization tandem mass spectrometry analysis (ESI-MS/MS), the *sn*-molecular species composition of PC, PE and PS was established (Figure 1C). We identified the main *sn*-molecular species of PC, PE, and PS to be 16:1/16:1, 16:0/16:1, 16:1/18:1, 16:0/18:1, and 18:0/16:1. With increasing culture temperature, PC mainly increases monounsaturated species 16:0/16:1 and 16:0/18:1 at the expense of 16:1/16:1, whereas PS and PE increase diunsaturated 16:1/18:1 at the expense of monounsaturated 16:0/16:1. Generally, upon increasing temperature, the acyl chain at the *sn*-1 position was found to decrease in unsaturation, whereas the acyl chain at the *sn*-2 position increases in length.

## Results

### Adaptation of lipid acyl chain composition to growth temperature

We first determined lipid acyl chain composition of wild type yeast cultured in synthetic defined glucose media at different temperatures by gas chromatography of fatty acid methyl esters obtained from total lipid extracts. As expected, the main acyl chains detected were C16:0, C16:1, C18:0 and C18:1 and comprise over 95% of total detected acyl chains (Figure 2A). In addition, low amounts of C12:0, C14:0 and C14:1 were detected (**Table S1**). When yeast is cultured at higher temperatures, lengthening of acyl chains is observed due to an increase in C18-acyl chains at the expense of C16:1 (Figure 2B). Interestingly, when cultured at 37°C acyl chains are both longer and more saturated compared to 30°C, whereas in cells cultured at 23°C acyl chains are shorter, while saturation is similar to 30°C. These observations are in agreement with previous reports (Martin, Oh, and Jiang 2007; Suutari, Liukkonen, and Laakso 1990; Hunter and Rose 1972).

**Figure 2.**
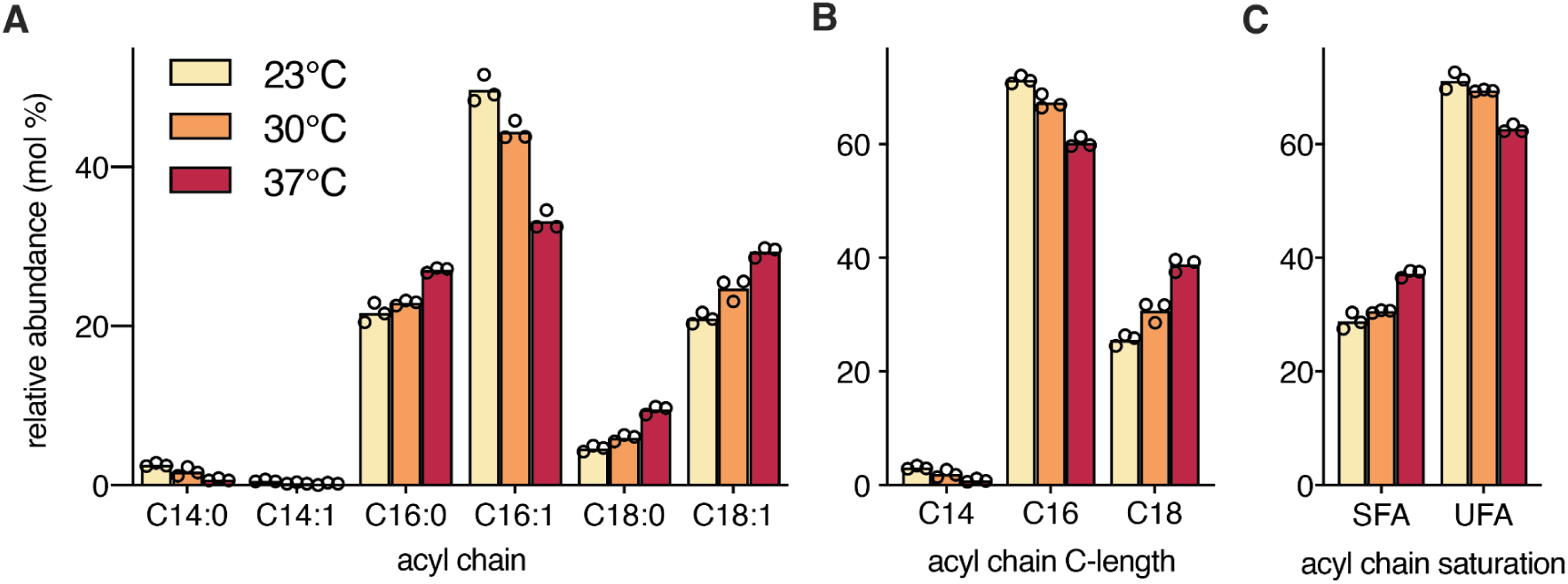
– Adaptation of the yeast total acyl chain composition to culture temperature. (A) Total acyl chain composition, (B) acyl chain length, and (C) content of saturated (SFA) and unsaturated acyl chains (UFA) of wild type yeast cultured to mid-log phase in SD medium at the indicated temperatures. Total lipids were extracted, and acyl chains were analyzed as methyl esters by GC-FID. Bars represent the mean of three biological replicate values shown as circles. (Data has previously been reported in (Renne and de Kroon 2018)). Underlying data for all sum molecular species can be found in Table S1.

### Adaptation of PC, PE, and PS lipid species at the sum-species level

Next, we analysed PC, PE, and PS at the sum-species (sum composition) level by traditional nESI-MS/MS using a triple quadrupole mass spectrometer. The sum-species profiles of PC, PE and PS consist mainly of 32:2, 32:1, 34:2 and 34:1 (Figure 3**, Table S2**), in agreement with previous reports (Klose et al. 2012; Schneiter et al. 1999; Casanovas et al. 2015). Notably, these four together comprise roughly 90% of the total PS, PE, and PC species, independent of the growth temperature (Figure 3).

**Figure 3.**
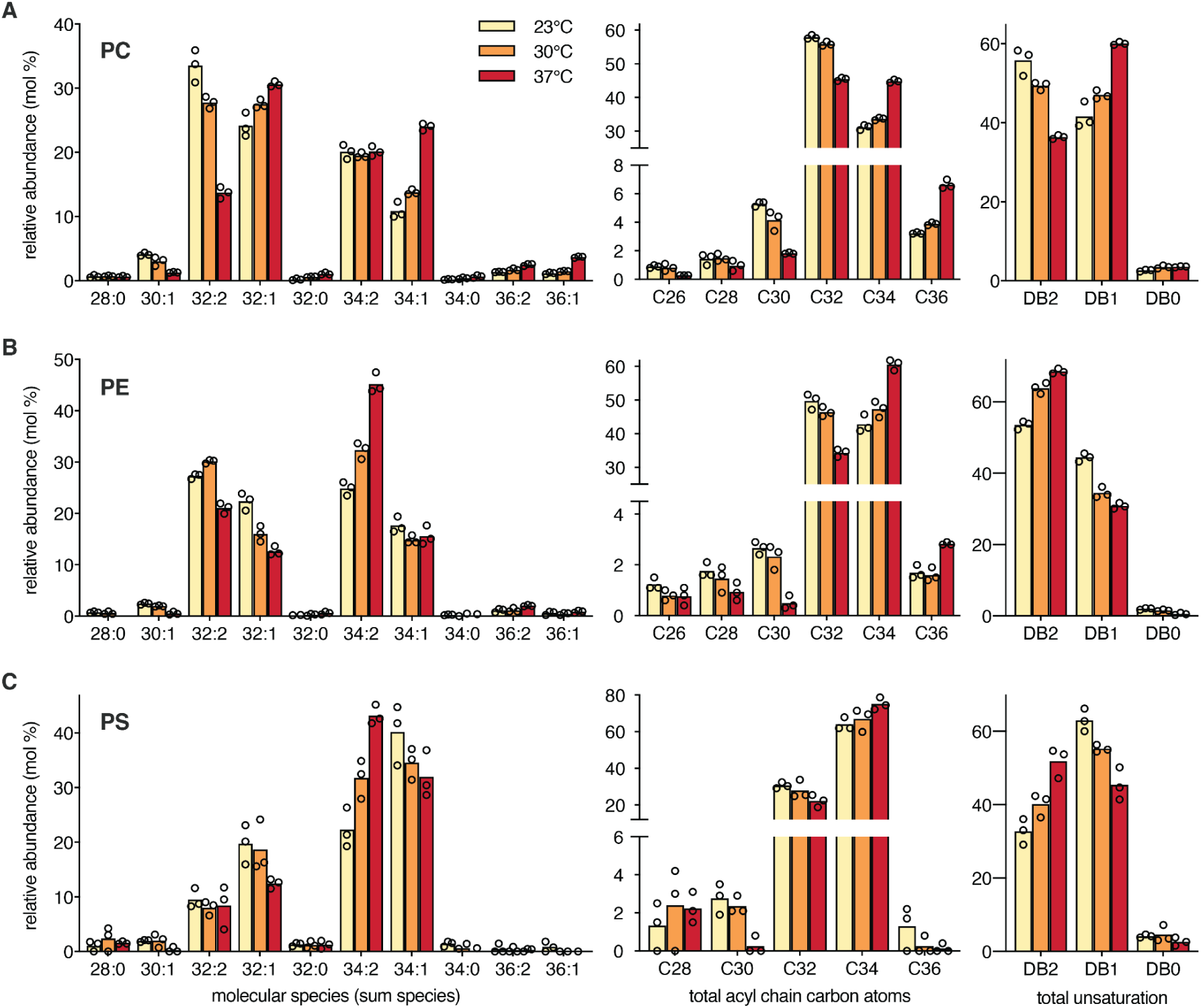
– Adaptation of phospholipid sum molecular species composition to culture temperature. Sum-species composition, total length, and number of double bonds of (A) PC, (B) PE, and (C) PS from wild type yeast cultured at the indicated temperatures, as determined by ESI-MS/MS. Bars represent the mean of three biological replicates, with individual values indicated by open circles. Total acyl chain length and number of double bonds are presented as the sum of the number of carbon atoms and the number of double bonds in both acyl chains, respectively. Underlying data for all sum molecular species can be found in Table S2.

At higher culture temperature, the PC sum-species composition increases in total acyl chain length and saturation (Figure 3A). These changes mainly result from a decrease in the relative abundance of PC32:2 and an increase in PC34:1. The relative abundance of PC32:1 varies only slightly and PC34:2 does not change. The PC species composition differs only slightly between 23°C and 30°C, whereas changes in composition are more drastic from 30°C to 37°C.

Compared to PC, the PE sum-species composition is enriched in 34:2 species at 30°C and 37°C, leading to higher overall PE unsaturation (Figure 3B). At 23°C, the relative abundance of di-unsaturated species is similar between PE and PC. The total acyl chain length of PE species increases at elevated culture temperature, mainly due to the higher PE34:2 abundance, at the expense of both PE32:2 and PE32:1 (Figure 3B). Furthermore, PE increases in total unsaturation with increasing temperature (Figure 3B), in contrast to PC where unsaturation decreases.

Compared to PE and PC, PS is strikingly enriched in C34 lipid species (Figure 3C). Furthermore, PS is enriched in mono-unsaturated species compared to PE. The adaptation of the PS sum-species composition to increasing culture temperature follows the same trends as PE with increasing acyl chain length and overall unsaturation. The PS composition shows a strong increase in 34:2 species with increasing temperature at the expense of 32:2, 32:1 and 34:1 (Figure 3C).

### Adaptation of the PC, PE, and PS sn-molecular species composition

To gain in-depth information on the lipid species adaptation at the level of fully defined molecular species structure, we deployed CID/OzID mass spectrometry (Brown, Mitchell, and Blanksby 2011; Pham et al. 2014; Maccarone et al. 2014). We obtained structural information for PC, PE, and PS and were able to identify 58, 34, and 22 different *sn*-molecular species (or positional isomers), respectively (Figure 4**, Table S3-5**). By superimposing the isomeric *sn*-molecular species distribution on the relative abundance of sum-species within each lipid class (**Table S2**), we determined the relative abundance of all *sn*-regioisomeric species within each lipid class (Figure 1B**, Table S6**).

**Figure 4.**
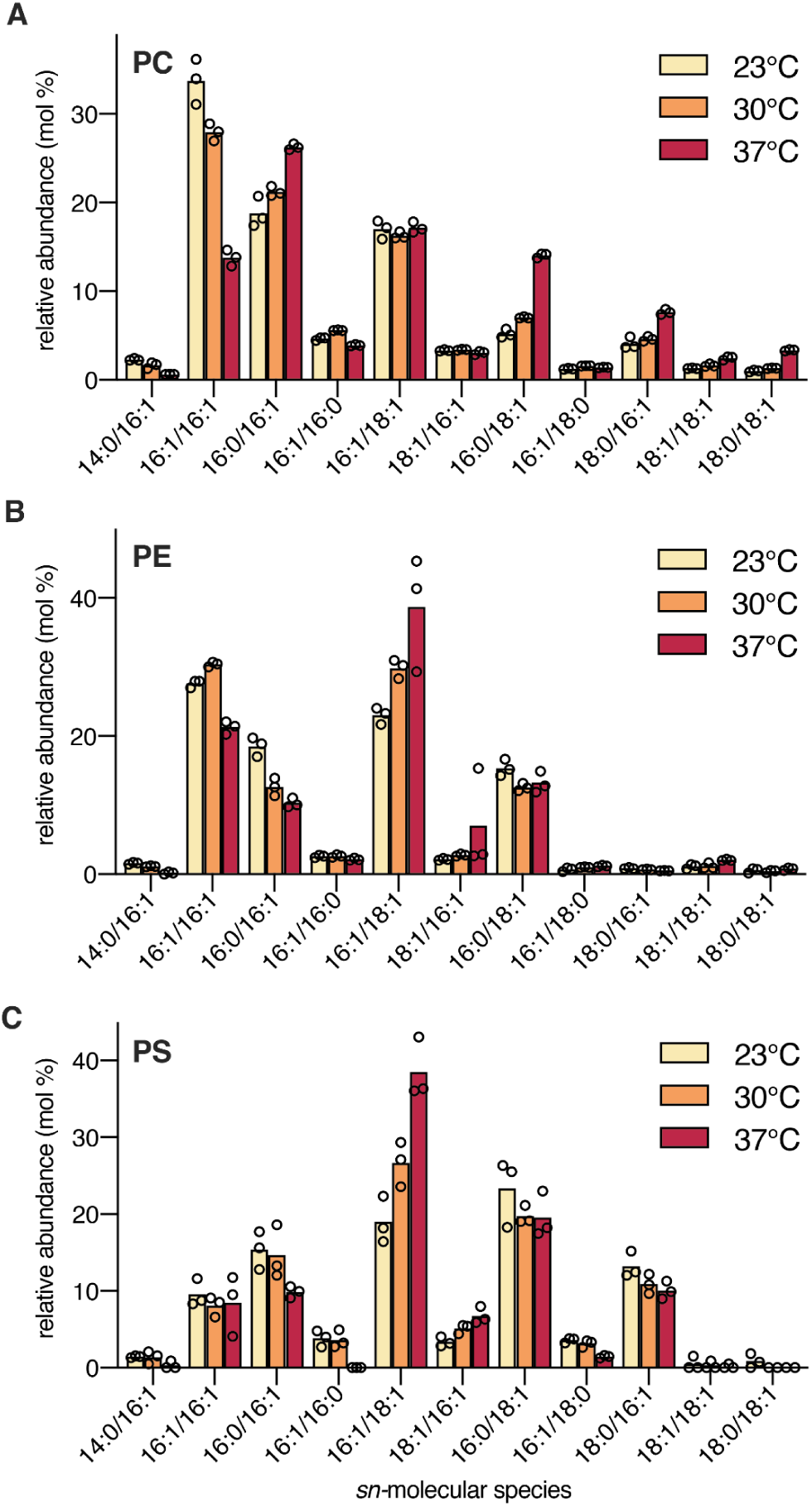
– Adaptation of phospholipid sn-molecular species composition to culture temperature. *sn*-molecular species composition of (A) PC, (B) PE, and (C) PS from wild type yeast cultured at the indicated temperatures. Within each sum-species, the abundance of different lipid isomers was determined by OzID mass spectrometry. sn-molecular species composition was calculated as described in the text. Bars represent the mean of three biological replicates, with individual values indicated by open circles. Underlying data for all *sn*-molecular species can be found in Table S6.

We first compared the *sn*-molecular species composition of PC, PE, and PS at optimal culture temperature (30°C). The main species of PC, PE and PS were found to be 16:1/16:1, 16:0/16:1, 16:1/18:1, 16:0/18:1 and 18:0/16:1 (Figure 4). PC and PE are enriched in 16:1/16:1 compared to PS (Figure 4A**,B**), in line with PE and PC being enriched in 32:2 sum species (Figure 3B). Similarly, PS and PE are enriched in 16:1/18:1 species, compared to PC, in line with their enrichment in 34:2 sum species (Figure 3B**,C**). Interestingly, some *sn*-molecular species were found to be specifically enriched or reduced in one specific lipid class. PC is enriched in 16:0/16:1, compared to PS and PE. The 16:0/18:1 species decreased in relative abundance from PS to PE to PC. PE was found to be almost completely devoid of 18:0/16:1 species, whereas these are relatively abundant in PS and PC (Figure 4).

Next, we investigated temperature adaptation of PC, PE and PS at the level of *sn*-molecular species. In agreement with the PC sum species adaptation to temperature (Figure 3A), PC16:1/16:1 decreases upon elevation of culture temperature, whereas PC 16:0/16:1, 16:0/18:1 and 18:0/16:1 increase (Figure 4A). The relative abundance of both PC 34:2 *sn-*regioisomers PC 16:1/18:1 and 18:1/16:1 did not change with altered growth temperature (Figure 4A), as was observed for PC 34:2 sum-species. The relative abundance of PC16:1/18:1 did not change with the temperature, whereas 16:1/18:1 species did increase in both PS and PE at higher temperatures. Inversely, whereas PC 16:0/18:1 increased with higher temperatures, 16:0/18:1 PS and PE are only barely altered. Moreover, the relative abundance of PC 16:1/16:0 and PC 16:1/18:0 hardly changed with temperature, even though other *sn*-molecular species within the corresponding sum species (PC32:1, and PC 34:1, respectively) did show variation.

Taken together, the data shows that the *sn*-molecular species composition of PC on one hand, *versus* PE and PS on the other, are altered in distinct manners upon variation of growth temperature.

### Adaptation at the sn-1 and sn-2 positions of PC, PE, and PS

The obtained *sn*-molecular species composition allows comparison of the PL acyl chain composition at the *sn-*1 and *sn*-2 position in relation to overall acyl chain composition per PL class (Figure 5). At 30°C, PC and PE are enriched in C16:1 acyl chains compared to PS (Figure 5). Compared to PS, the C16:1 enrichment is mainly at the expense of C16:0 and C18:0 in PE, and mainly at the expense of mainly C18:1 in PC. Similar trends have been observed previously (Ejsing *et al*, 2009; Wagner & Paltauf, 1994).

**Figure 5.**
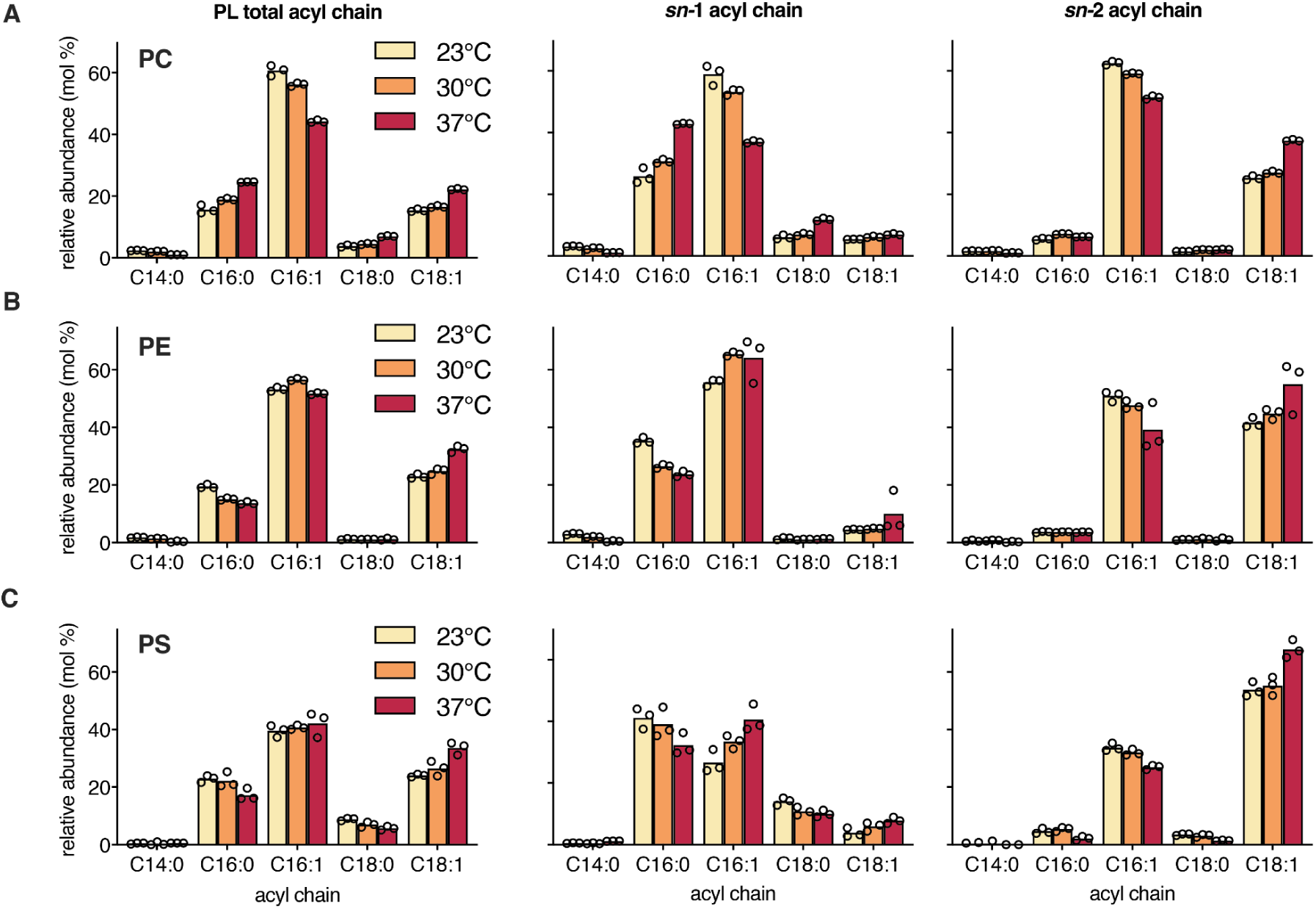
– Adaptation of phospholipid acyl chain composition to culture temperature. Total acyl chain composition, *sn*-1 acyl chain composition, and *sn*-2 acyl chain composition of (A) PC, (B) PE, and (C) PS from wild type yeast cultured at indicated temperatures. Bars represent the mean of three biological replicates, with individual values indicated by open circles.

At higher temperature, the abundance of C16:0, C18:0 and C18:1 in PC increases at the expense of C16:1 (Figure 5A), similar to the trends observed for the total acyl chain composition (Figure 2). Interestingly, the acyl chain compositions of PE and PS are altered to a less extent compared to PC. Both PE and PS have increased C18:1 at 37°C, at the expense of C16:0 in PE and C16:0/C18:0 in PS (Figure 5B**,C**). Overall, the increase of C18 at the expense of C16, is less pronounced in PS and PE, compared to PC.

Next, we compared acyl chain compositions at the *sn-*1 and *sn*-2 positions of the different phospholipids. At 30°C, the *sn-*1 position of PS, PE and PC is strongly enriched in saturated acyl chains, whereas the *sn*-2 position is enriched in C18:1 (Figure 5) as reported previously (Wagner & Paltauf, 1994). The difference between the *sn*-1 and *sn*-2 acyl chain composition is most pronounced in PE, with the *sn*-1 position containing almost exclusively C16:0 and C16:1, whereas at the *sn*-2 mainly C16:1 and C18:1 are present (Figure 5B). At the PC *sn-*2 position C16:1 is more abundant than C18:1, yet in PS the opposite is observed (Figure 5B). Interestingly, PS and PC have a much higher C18:0 content compared to PE (Figure 5).

Comparison of the *sn*-1 and *sn*-2 acyl chain composition at different culture temperatures revealed distinct adaptations to culture temperature at the *sn*-1 and *sn*-2 position. In PC, C16:0 increases at the *sn*-1 position with increasing temperature, at the expense of C16:1, whereas the proportion of C16:0 at the *sn*-1 position in PS and PE shows a slight drop (Figure 5). At the *sn*-2 position, increasing culture temperature leads to an increase in C18:1 abundance, at the expense of C16:1, similar in PC, PE, PS. These observations indicate that temperature adaptation at the *sn*-1 position mainly occurs via alteration of acyl chain unsaturation (C16:0/C16:1), as opposed to alteration of acyl chain length (C18:1/C16:1) at the *sn*-2 position.

## Discussion

Yeast adapts its lipid composition in response to altered culture temperature, to maintain homeostasis of membrane physical properties. Recent studies have employed high resolution mass spectrometry-based lipidomics to describe the yeast lipidome at the level of sum composition (Klose et al. 2012) and molecular composition (Ejsing et al. 2009), however to our knowledge analysis of the yeast lipidome at the full molecular structure level has not been reported. Taking advantage of CID/OzID mass spectrometry, the present study analyses the yeast glycerophospholipid species composition at the level of *sn*-regioisomers. We identified 58, 34, and 22 different molecular species of PC, PE, and PS respectively, adding an extra layer of information to the yeast lipidome that may shed increased light on lipid metabolism and metabolic relationships.

The main molecular species of PC, PE and PS were found to be 16:0/16:1, 16:0/18:1, 16:1/16:1, 16:1/18:1, and 18:0/16:1 (Figure 4), in agreement with the major sum species reported to be 32:2, 32:1, 34:2 and 34:1 (Klose et al. 2012; Ejsing et al. 2009). Notably, we observed a clear preference for the *sn*-regioisomer species of 32:1 and 34:1 species. While both 16:0/16:1 and 16:0/18:1 are found as bulk PL species, the corresponding *sn*-regioisomers 16:1/16:0 and 18:1/16:0 are much less abundant (Figure 3). In fact, 18:1/16:0 isomers are only found in trace amounts. This indicates a preference for the synthesis of PLs with a saturated C16:0 acyl chain at the *sn*-1 position, and an unsaturated acyl chain at the *sn*-2 position (Figure 4), in agreement with previous work (Wagner and Paltauf 1994).

In agreement with previous reports, the total acyl chain composition varies drastically depending on the culture temperature (Figure 2) (Martin, Oh, and Jiang 2007; Suutari, Liukkonen, and Laakso 1990). Interestingly, when compared to the optimal temperature (30°C), the acyl chain composition at lower temperatures (23°C) shows only small variation, whereas the acyl chain composition at higher temperature (37°C) differed more dramatically (Figure 2). In general, with increasing temperatures the acyl chain length increases mainly by an enrichment of C18 acyl chains at the expense of C16 (Figure 2**, 5**), causing an increase in the species length of PC, PE, and PS (Figure 3). The length of the acyl chains is determined by fatty acid synthesis (by the FAS complex) and acyl-CoA elongation (by Elo1/2/3p) (Martin, Oh, and Jiang 2007; de Kroon, Rijken, and De Smet 2013; Renne and de Kroon 2018). Consistent with our observations, temperature-dependence of the fatty acid length synthesised by the FAS complex has been observed *in vitro* (Okuyama et al. 1979). At elevated temperature (37°C) the total acyl chain composition shows an increase in acyl chain saturation (Figure 2), conforming to the previously reported temperature dependent regulation of the expression of the acyl-CoA desaturates Ole1p (Nakagawa et al. 2002). The increase in total UFA is due to an increase in C18:1 but not C16:1 (Figure 2), in line with C18:1 being the preferred product of Ole1p (De Smet et al. 2012). Consistent with the total acyl chain composition, at elevated temperature the abundance of C18:1 increases in PC, PE and PS (Figure 5).

Notably, the temperature-adaptation of the PC *sn*-regioisomer composition is markedly different from that of PS and PE (Figure 3), showing increases in PC 16:0/16:1 and PC 18:0/16:1 and a decrease in PC 16:1/16:1 upon elevated temperature. Furthermore, 16:1/18:1 species increase in both PS and PE when culture temperature increases, whereas the PC 16:1/18:1 level is similar at all temperatures yet PC 16:0/18:1 increases. Under the culture conditions used, *i.e.* using media devoid of choline and ethanolamine, net synthesis of PC depends on the PE methylation route, which in turn relies on PS decarboxylation (de Kroon 2007). The conversion of PS to PE as well as PE-methylation yielding PC preferentially produce diunsaturated species (Boumann et al. 2003; Renne et al. 2022), suggesting that the temperature-induced increase in PC species with a saturated acyl chain on the *sn*-1 position and a unsaturated acyl chain at *sn*-2 (in example PC 16:0/16:1) does not originate from *de novo* synthesis. Instead, these mono unsaturated PC species could be formed after turnover of PC and recycling of the choline headgroups via the CDP-choline route (or Kennedy pathway) (Fernández-Murray and McMaster 2007) and/or via an acyl chain remodelling process (Renne et al. 2015; Patton-Vogt and de Kroon 2020). Indeed, PC turnover, including PC deacylation to yield glycerophosphocholine, and PC synthesis via the CDP-choline route are increased at 37°C (Dowd, Bier, and Patton-Vogt 2001). Furthermore, increasing PC saturation by acyl chain remodelling occurs mainly at the *sn*-1 position (Boumann et al. 2003), and likely depends on de-acylation by the phospholipase B Plb1 and re-acylation by Gpc1p (Anaokar et al. 2019; De Smet et al. 2013).

The data in this study further expand the knowledge on the changes in the lipidome in response to altered growth temperature. The homeoviscous adaptation model posits that these changes are required to maintain membrane physical properties, such as membrane fluidity (Hazel 1995). Indeed, *in vitro* biophysical studies utilising model membranes have shown how changes in the lipid composition affect the physical properties of a membrane. However, for complex lipid compositions, such as those observed in biological membranes, it is difficult to interpret how changes in the abundance of certain lipids affect the physical properties of bulk membranes. This is complicated even further by the fact that the chemico-physical properties have only been determined (experimentally) for a relatively small amount of lipid species (Renne and de Kroon 2018; Koynova and Caffrey 1994). Recent studies have extrapolated data from trends observed in experimental studies (Reinhard et al. 2023) or utilised atomistic molecular dynamics simulations (Smith et al. 2021) to bypass this problem, but the lack of experimental studies addressing the chemico-physical properties of different lipids remains. By combining synthetic organic chemistry for the synthesis of specific lipid species and biophysical studies to determine lipid physico-chemical properties, future studies should greatly expand our knowledge of lipid properties. These studies, in conjunction with state-of-the-art lipidomics workflows, will be beneficial to gaining mechanistic understanding of how changes in the lipid composition affect membrane biophysics.

## Materials and Methods

### Yeast strain and culture conditions

Wild type yeast (BY4741; from EuroSCARF) was cultured in synthetic complete dextrose (SD) medium, containing per liter: 6.7 g yeast nitrogen base with ammonium sulfate (Difco, B/D Bioscience), 20 mg adenine, 20 mg arginine, 20 mg histidine, 60 mg leucine, 230 mg lysine, 20 mg methionine, 300 mg threonine, 20 mg tryptophan, 40 mg uracil, and 20 g glucose. All components were added to autoclaved MQ water from filter sterilized stock solutions.

Single colonies were pre-cultured for 24 h in SD medium at 30°C, inoculated in 100 mL pre-warmed SD medium (23°C, 30°C, 37°C) at OD_600_ < 0.02, and cultured at the indicated temperature to mid-exponential growth (OD_600_ ≈ 1). All cultures were grown while shaking at 200 rpm in tubes (pre-cultures) or Erlenmeyer-flasks with a ratio of culture volume to tube/flask volume of at most 1:3. Culture density was determined by measuring the optical density at 600 nm (OD_600_) on a Novaspec II single beam spectrophotometer (Pharmacia Biotech). Cells were harvested by centrifugation (3000 g, 4 min, 4°C), washed with MQ water and stored at -20°C. Lipids were extracted as described below, and individual extracts were subjected to lipid analysis by both GC and MS approaches.

### Cell disruption and lipid extraction

An aliquot of cells corresponding to 40 – 60 OD_600_ units was resuspended in a final volume of approx. 600 μL MQ and added to a 2 mL Eppendorf tube containing 0.5 mL acid washed glass beads (Sigma Aldrich). Cells were disrupted by bead bashing for 5 min using a Vortex Genie II with a Turbomix attachment (Scientific Industries). Tubes were kept on ice between all steps. Lipids were extracted using a modified version of the lipid extraction procedure described by Bligh and Dyer (Bligh and Dyer 1959), as previously described (Renne et al. 2022). Briefly, the 600 μL cell lysate was added to a glass tube containing 1360 μL MeOH, 620 μL CHCl_3_, and 20 μL 1 M HCl was added. The tube was vortexed briefly and kept on ice for at least 15 min. 600 μL CHCl_3_ and 600 μL 0.1 M HCl were added to induce phase separation. The tube was vortexed briefly, kept on ice for 2 min, centrifuged for 4 min at 3000 g at 4°C, and the organic phase was transferred to a new glass tube. Next, 120 μL 1 M KCl was added to the water phase, the water phase was washed with 600 μL CHCl_3_, and the organic phases were pooled. The combined organic phase was washed with 0.1 M KCl, transferred to a new glass tube, and after adding 120 μL isopropanol dried in a water bath (40°C) under N_2_-flow. After dissolving the lipid film in 500 μL CHCl_3_, the phospholipid concentration was determined using the inorganic phosphate determination as described by Rouser *et al*. (Rouser, Fleischer, and Yamamoto 1970).

### Analysis of total acyl chain composition by GC-FID

The total acyl chain composition was determined by gas chromatography analysis of fatty ester methyl esters as described previously (Renne et al. 2022). Total lipid extracts corresponding to 100 nmol phospholipid phosphorus were transesterified by heating for 2.5 h at 70°C in 2.5 mL 2.5% (v/v) methanolic H_2_SO_4_. Fatty acid methyl esters (FAMEs) were extracted by addition of 2.5 mL hexane and 2.5 mL water. After collecting the organic phase, the aqueous phase was washed with hexane, the organic phases were pooled and washed with water until the pH of the water phase was neutral. 120 μL isopropanol was added and the sample was dried in a water bath (40°C) under N_2_-flow. The FAME were dissolved in 1 mL hexane, transferred to an Eppendorf tube and centrifuged for 10 min (14,000 g at 4°C) to remove any residual particles. Next, 900 μL of the supernatant was concentrated to a volume of 50 - 100 μL under N_2_-flow.

FAMEs were analysed by GC-FID using splitless injection (2 μL injection volume, inlet temperature 230°C, 1 min splitless time with a flow of 20 mL/min) on a Trace*GC Ultra* (Thermo Scientific) equipped with a biscyanopropyl-polysiloxane column (Restek), using N_2_ as carrier gas (1.3 mL/min, constant flow). After injection, the samples were concentrated on the column by keeping the temperature at 40°C for 1 min, after which the column was rapidly heated to 160°C (30°C/min). FAMEs were separated using a temperature gradient from 160°C to 220°C (4°C/min) and signal was detected using FID at 250°C (H_2_/air with N_2_ as make-up). Integrated peaks were identified and calibrated using a commercially available FAME standard (63-B, Nu-Chek-Prep). Acyl chain compositions are presented as mol% of the four most abundant acyl chains (C16:0, C16:1, C18:0 and C18:1) that were recovered in all samples.

### Electrospray ionisation - tandem mass spectrometry analysis

Analysis of lipids at the sum-species levels was carried out using a chip-based nano-electrospray ionisation source (TriVersa NanoMate, Advion, Ithaca, NY, USA) coupled to a QTRAP 5500 triple quadrupole mass spectrometer (SCIEX, Framingham, MA, USA) (Norris et al. 2015). Dried lipid extracts (100 nmol phospholipid phosphorus) were dissolved in 100 µL of CH_2_Cl_2_ containing 0.01% butylhydroxytoluene (BHT; SigmaAldrich, NSW, Australia) to yield a solution *ca.* 1 mM total phospholipid concentration. Solutions were further diluted to 1 µM using 2:1 MeOH:CHCl_3_ (ThermoFisher Scientific, Australia) with 5 mM ammonium acetate (SigmaAldrich, NSW, Australia), then loaded into an Eppendorf twin-tec PCR 96 well plate (Eppendorf, Hamburg, Germany). A clean, conductive pipette tip in the NanoMate collected 10 µL of lipid solution from the 96 well plate maintained at 10 °C and delivered it to the mass spectrometer via a nano ESI chip (4.1 µm nozzle diameter). Nitrogen was delivered at 0.4 psi to the pipette tip while the spray voltage was set to 1.15 kV for both polarities. All lipid classes targeted for profiling are listed in **Table 1**. For each of positive and negative modes, typical mass spectrometer conditions included a declustering potential (DP) of 100 V, entrance potential (EP) of 10 V and a scan rate of 200 *m/z* per second. Data was acquired using Analyst v1.6.2 software. Lipid profiling was performed using LipidView v1.2 (Sciex) software, including smoothing, identification (sum composition), removal of isotope contributions and correction for isotopic distributions. No absolute quantification of lipid abundance was carried out.

**Table 1.** - List of scans carried out for sum composition lipid profiling on a Sciex Q-Trap 5500. Abbreviations used: Scan types: PIS, precursor ion scan; NLS, neutral loss scan;. Listed Collision Energies (CE) and Collision Cell Exit Potentials (CXP) are the values programmed into the Analyst control software (version 1.6.2).

| Lipid | Polarity | FA | Scan type | Mass | CE | Scanned range<br>(m/z) | CXP |
| --- | --- | --- | --- | --- | --- | --- | --- |
| PC | + |  | PIS | 184.1 | 40 | 490-820 | 8 |
| PE | + |  | NLS | 141.0 | 30 | 440-770 | 8 |
| PS | + |  | PIS | 185.0 | 25 | 480-820 | 8 |
|  | - |  | NLS | 87.0 | 30 | 480-820 | 11 |
| FA | - | 8:0 | PIS | 143.1 | 55 | 400-750 | 11 |
|  | - | 8:1 | PIS | 141.1 | 55 | 400-750 | 11 |
|  | - | 10:0 | PIS | 171.1 | 55 | 420-780 | 11 |
|  | - | 10:1 | PIS | 169.1 | 55 | 420-780 | 11 |
|  | - | 12:0 | PIS | 199.2 | 55 | 450-810 | 11 |
|  | - | 12:1 | PIS | 197.2 | 55 | 450-810 | 11 |
|  | - | 14:0 | PIS | 227.2 | 55 | 480-840 | 11 |
|  | - | 14:1 | PIS | 225.2 | 55 | 480-840 | 11 |
|  | - | 16:0 | PIS | 255.2 | 55 | 510-870 | 11 |
|  | - | 16:1 | PIS | 253.2 | 55 | 510-870 | 11 |
|  | - | 18:0 | PIS | 283.3 | 55 | 540-890 | 11 |
|  | - | 18:1 | PIS | 281.2 | 55 | 540-890 | 11 |

### Advanced structural characterisation by CID/OzID mass spectrometry

Advanced structural characterisation was performed using a modified Orbitrap Fusion™ Tribrid™ mass spectrometer (ThermoFisher, San Jose, USA) coupled to a high concentration ozone generator (see below). Each lipid extract (10 µM in 2:1 MeOH:CHCl_3_ with 100 µM sodium acetate) was directly infused via syringe pump at 7 µL/min through an Easy-Max NG H-ESI source equipped with the high-flow ESI capillary (Thermo part number THI80000-60317). For each targeted phospholipid subclass an Xcalibur method containing three successive tandem MS experiments was deployed: negative mode CID (fatty acyl sum composition) and positive mode CID/OzID (fatty acyl *sn*-position) for 1, 3, and 9 minutes acquisition, respectively. A total of 39 target ions were selected for each class (PC, PS and PE) and subjected to each of the CID and CID/OzID characterisation methods. Targets were selected to include all possible combinations for the three listed phospholipid subclasses bearing fatty acyl chains containing 8 to 20 carbons with none or one carbon-carbon double bond. The following parameters were common to all studied phospholipid subclasses and tandem MS methods: Orbitrap scans were acquired at 120,000 resolving power (FWHM *m/z* 200) between 200 and 1000 m/z. The quadrupole was used to isolate precursor ions using a 1 *m/z* width (0.7 *m/z* for CID/OzID on PC). Ion injection times were 247.5 ms and the ion transfer tube and vaporizer temperature were 300 and 40 °C, respectively. The RF lens potential was set to 60% except for the PC CID/OzID and OzID scans where it was 65%. For PC and PE subclasses a spray voltage of (-)3.0/(+)3.9 kV was used whereas it was (-)3.0/(+)4.7 for PS. Nitrogen was used as the sheath, auxiliary and sweep gases which were respectively 5, 2.5 and 1 (PC); 5, 6 and 1 (PE); and 10, 10 and zero (PS). Negative mode CID experiments deployed normalised collision energy (CE) percentages of 27 (PC) and 28 (PE and PS) with a 10 ms activation time. CID/OzID experiments were conducted in the positive ion mode employing normalised HCD collision energy percentages of 25 (PC and PS) and 30 (PE). Normalised CE percentages were set to zero during 150 ms activation time in the ion trap where an isolation width of 2 Th was programmed.

### Generation of ozone for CID/OzID

CID/OzID was performed on an Orbitrap Fusion mass spectrometer modified to allow (*i*) introduction of ozone into the Linear Ion Trap (LIT) region and (*ii*) increased LIT isolation time up to 4 seconds. The hardware modifications to deliver ozone to the LIT were modelled after those reported by Paine *et al*. (Paine et al. 2018). High purity oxygen with 0.5% nitrogen (Coregas, NSW, Australia) was fed at 20 psig and 200 mL/min through an ozone generator (Titan 30, Absolute Systems, Edmonton, AB, Canada) set to enable ca. 15% (weight % O_2_) ozone production. The flow was directed through a catalytic destruct (INUSA, MA, USA) with resulting oxygen directed outside the building. The ozone generator was interlocked to an ambient ozone monitor (106-L, 2B Tech., CO, USA) in the event that room ozone levels exceeded 60 ppb. Prior to ozone destruction, part of the gas flow was diverted through a PEEKsil restriction (1/16” OD x 25 micron ID, length 10 cm) and then combined with the helium flow into the linear ion trap. Helium flow was regulated by both a regulator (0-50 psi, Swagelok, Australia) and PEEKsil restriction (1/16” OD x 50 micron ID, length 10 cm). The regulator was adjusted in the absence of ozone delivery to the mass spectrometer to achieve a linear ion trap pressure specified for normal operation. A software patch was installed on the Orbitrap Fusion control PC running FusionTune 3.0 to allow linear ion trap isolation times up to 4 seconds by programming the Scan Description field in the Method Editor section of Xcalibur (version 4.0).

### Data Analysis

Data analysis was automated using an in-house python script (Python 3.6.5) for each experiment type and full characterisation was obtained where possible by cross reference of fragments obtained for each experiment.

Mass spectra were converted to *m/z* versus ion counts, intensities were extracted to a spreadsheet for any mass spectral peak with greater than 500 counts, and corresponding *m/z* values were compared within 50 mDa to one of three look-up tables depending on the experiment type to yield lipid assignments with corresponding ion counts. Assignments were cross-referenced manually between the different MS analyses to provide the final lipid identification.

### Author contributions

M.F.R. conceived and designed the study. C.K., A.T.M., and M.F.R. conducted experiments. All authors analysed data, interpreted data, and discussed results. M.F.R. wrote the manuscript with input from the other authors. All authors reviewed and edited the manuscript.

## Acknowledgements

The authors acknowledge use of the University of Wollongong Mass Spectrometry Facility (MSF). We thank Christer Ejsing, David Marshall, and Antoinette Killian for valuable discussions, and Juan Dominguez-Pardo, Hanaa Hariri, and Robert Ernst for critical reading of the manuscript. UOW researchers would like to thank David Marshall, Berwyck Poad, and Stephen Blanksby from QUT for assistance modifying hardware for Ozone-Induced Dissociation experiments. In addition, we thank Edward Gonzalez, Shannon Eliuk, and Steve Binos at Thermo-Fisher Scientific for providing the FusionTune software patch to extend the isolation time in the LIT of the Orbitrap Fusion mass spectrometer.

This research was supported by the Division of Chemical Sciences in the Netherlands, with financial aid from The Netherlands Organization for Scientific Research (to MFR) and a Summer Fellowship from the Federation of European Biochemical Societies (to MFR).

## List of supplementary information

**Table S1 - total acyl chain composition determined as fatty acid methyl esters by GC-FID**

Data is given as mean and standard deviation (SD) of 3 biological replicates.

**Table S2 - sum species composition of PC, PE, and PS as determined by ESI-MS/MS.**

Data is given as mean and standard deviation (SD) of 3 biological replicates.

**Table S3/S4/S5 - *sn*-regioisomer composition per sum species of PC (Table S3), PE (Table S4), and PS (Table S5) as determined by CID and CID/OzID FT-MS^n^.**

Data is given as mean and standard deviation (SD) of 3 biological replicates.

**Table S6 - *sn*-regioisomer composition of PC, PE and PS.**

Data is given as mean and standard deviation (SD) of 3 biological replicates. Values were determined by superimposing the *sn*-regioisomer composition (Table S3/S4/S5) on the sum species composition (Table S2).

